# Maternal progesterone and adipose mPRε in pregnancy regulate the embryonic nutritional state

**DOI:** 10.1101/2024.08.26.609823

**Authors:** Keita Watanabe, Mayu Yamano, Junki Miyamoto, Ryuji Ohue-Kitano, Yuki Masujima, Daiki Sasahara, Yuki Mouri, Nozomu Kono, Shunsuke Inuki, Fumitaka Osakada, Kentaro Nagaoka, Junken Aoki, Yuki Sugiura, Hiroaki Ohno, Eiji Kondoh, Ikuo Kimura

## Abstract

Sex steroid hormones such as progesterone play a pivotal role in reproductive functions and maintaining pregnancy; however, the impact of progesterone on the interaction between mother and embryo is unclear. Here, we demonstrate that the relationship between maternal progesterone and membrane progesterone receptor epsilon (mPRε) in adipose tissue regulates embryonic nutritional environment and growth after birth in mice. The activation of adipose mPRε by increased progesterone during pregnancy enhanced maternal insulin resistance through the production of prostaglandins, thereby efficiently providing glucose to embryos. The offspring of mPRε-deficient mothers exhibited metabolic dysfunction, whereas mPRε-deficient mothers with high-fat-diet-induced obesity exhibited improved insulin sensitivity. These findings establish the importance of progesterone as a nutritional regulator between mother and embryo, and suggest that mPRε modulators could be developed to treat pregnant glycemic control disorders such as gestational diabetes mellitus, as well as metabolic syndrome in offspring.

**Graphical Abstract:** 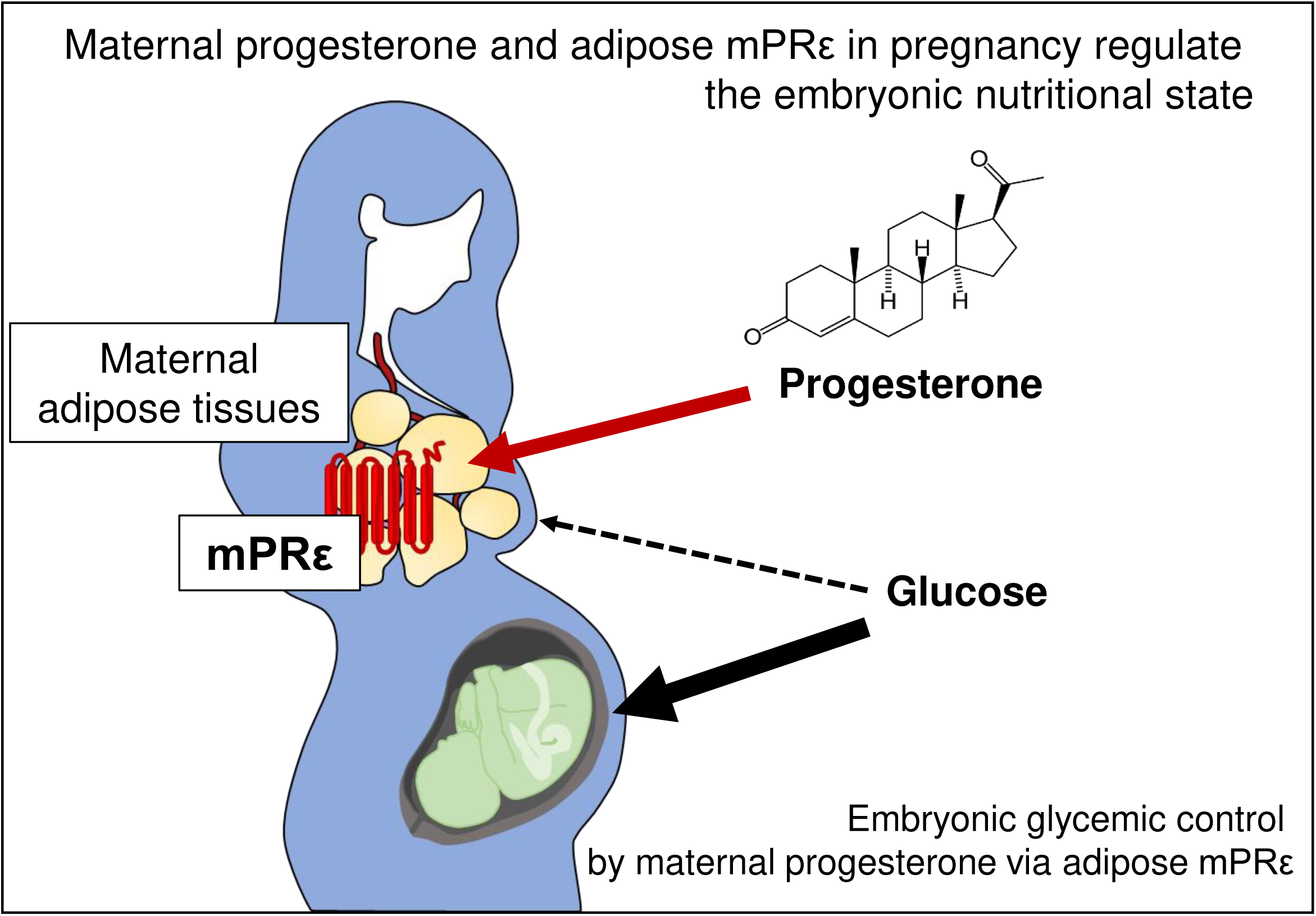

## Introduction

During the perinatal period, fetal growth is influenced by various maternal humoral factors, which directly impact the development of conditions such as allergies, neurological disorders, obesity, and diabetes in offspring^1–5^. For example, obesity-induced dysfunction of blood glucose control during pregnancy, gestational diabetes mellitus^6^, defined as glucose intolerance of variable degree caused by maternal overnutrition, and starvation owing to maternal malnutrition^1–3^ all lead to abnormal weights according to gestational age and pose a risk of excessive body weight gain and metabolic syndrome later in life^7–12^. Among maternal humoral factors, sex steroid hormones such as estrogen and progesterone are the most important for reproduction^13–15^. The peak levels of these hormones during pregnancy are over 10-fold higher than those during the estrus cycle^15,16^. Therefore, maternal sex steroid hormones are considered crucial not only for maintaining pregnancy in the mother but also for embryonic development. However, the underlying mechanisms and biological significance of maternal–embryonic crosstalk via maternal sex steroid hormones remain unclear.

In addition to reproductive functions, sex steroid hormones are involved in many physiological functions, including maternal behavior, aggression, emotion, feeding, circadian rhythm, sleep, and higher brain functions such as memory and learning^13–16^. Most of these hormones generate immediate responses that cannot be explained by nuclear receptors, and the mechanisms are unclear^17,18^. Instead, membrane progesterone receptors (mPRs) and GPR30, which are the respective membrane receptors for progesterone and estrogen^19,20^, may mediate the immediate physiological responses of sex steroid hormones, such as the activation of MAPK signaling, intracellular Ca^2+^ increase, and release of fatty acids from membrane phospholipids^17–21^.

mPRε/Paqr9 belongs to the progestin and AdipoQ receptor (PAQR) family, which includes five unique mPR families: mPRα/Paqr7, mPRβ/Paqr8, mPRγ/Paqr5, mPRδ/Paqr6, and mPRε^22–24^. mPRε can sense and respond to progesterone with EC_50_ values of approximately 13 nM^25,26^. Moreover, the progesterone–mPR signal is involved in GPCR signaling and the MAPK cascade^27^. Although the physiological functions of hepatic mPRε include the regulation of fasting-induced ketogenesis and fatty acid oxidation in the liver^28^, and those of pancreatic mPRε include the regulation of glucose and lipid homeostasis in diabetic mice^29^, the relationship between these phenotypes and progesterone as a ligand for mPRε remain unelucidated.

Considering the high expression of mPRε in white adipose tissues and high levels of progesterone in pregnancy, we aimed to clarify the interaction between progesterone and adipose mPRε during pregnancy and elucidate the mechanisms by which sex steroid hormones modulate the interaction between embryos and maternal metabolism using mPRε-deficient mice.

## Results

### *mPRε* is abundantly expressed in the adipose tissues

First, we examined mPRε expression in mice on postnatal day 49 (P49), during sexual maturation, using real-time quantitative PCR. *mPR*ε mRNA was detected in the liver, kidney, white adipose tissue (WAT), and brown adipose tissue in both males and females. Moreover, *mPR*ε expression significantly increased in the WATs of mice fed a high-fat diet (HFD) compared with those on normal chow (NC) (Fig. 1a). Regarding subtypes of the mPR family, only *mPR*ε was expressed in the WATs of both male and female mice (Fig. 1b). In these tissues, *mPR*ε was expressed in the mature adipocytes but not in the stromal vascular fraction (SVF), and expression was significantly higher under HFD feeding than that under NC feeding (Fig. 1c). Moreover, *mPRε* expression gradually increased with adipogenesis of mouse embryonic fibroblasts (MEFs) in both males and females (Fig. 1d). Notably, nuclear progesterone receptor expression was only observed in the SVF of female WATs (Fig. 1e). Thus, our findings suggest that *mPRε* is expressed in the adipocytes of WATs in both male and female mice.

**Fig. 1.**
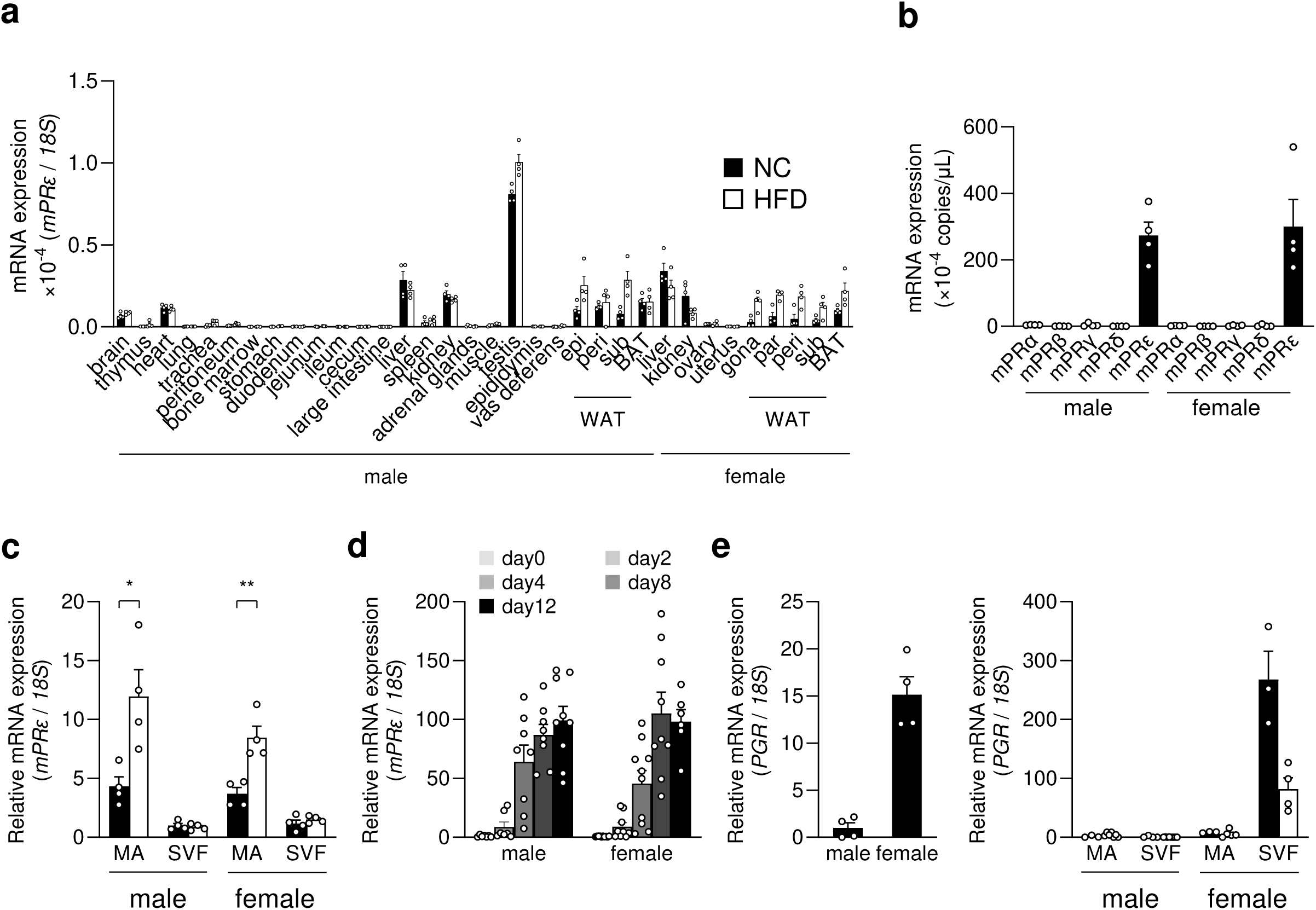
*mPRε* is sufficiently expressed in white adipose tissues. (a) *mPRε* mRNA expression in mouse tissues (Postnatal day 49: P49) measured by quantitative RT-PCR (n = 3–4). WATs: White adipose tissues (epididymal adipose tissue); BAT: Brown adipose tissue. Control *18S* rRNA expression. Statistical analysis was performed using Student’s t-test. NC, normal chow; HFD, high-fat diet. (b) Expression of mPR family members in mouse WAT (n = 4) (postnatal day 49, P49). (c) mPRε expression in mature adipocytes (MA) and a stromal vascular fraction (SVF) of mice fed a normal or high-fat diet (n = 4). (d) *mPRε* mRNA expression in MEF-derived adipocytes during adipogenesis (n = 6–10). (e) Expression of the progesterone nuclear receptor PGR in mouse WATs (left). PGR expression in the MA and SVF of mice fed NC or HFD (n = 3–8). (Post-natal day 49: P49). **P < 0.01, *P < 0.05 (Student’s t-test). Results are presented as the mean ± standard error of the mean (SE).

### Progesterone exacerbates insulin resistance via mPRε

Next, we generated mPRε-deficient mice to clarify the physiological function of mPRε on energy metabolism (Extended Data Fig. 1a–c). In the HFD-induced obese mouse model, the body weights of mPRε-deficient mice did not differ significantly from those of wild-type mice in either males or females during growth (Fig. 2a). Similarly, the tissue weights and metabolic parameters at 16 weeks of age were comparable between wild-type and mPRε-deficient mice for both males (Extended Data Fig. 2a–c) and females (Extended Data Fig. 2d–f). Plasma progesterone levels were less than 10 nM, (< EC_50_ of mPRε) even in females (Fig. 2b) because the corpus luteum degenerates after ovulation in mice unless certain hormones are released^30^. Therefore, we externally stimulated mPRε by subcutaneously (s.c.) administering progesterone. Progesterone administration (s.c.) increased plasma progesterone levels (∼50 nM) in both males and females of both wild-type and mPRε-deficient mice (Extended Data Fig. 3a). According to glucose and insulin tolerance tests, progesterone administration (s.c.) exacerbated insulin resistance in both male and female wild-type mice; however, this effect was abolished in both male and female mPRε-deficient mice (Fig. 2c–f). Progesterone administration did not lead to significant differences in plasma insulin levels between wild-type and mPRε-deficient mice (Extended Data Fig. 3b).

**Fig. 2.**
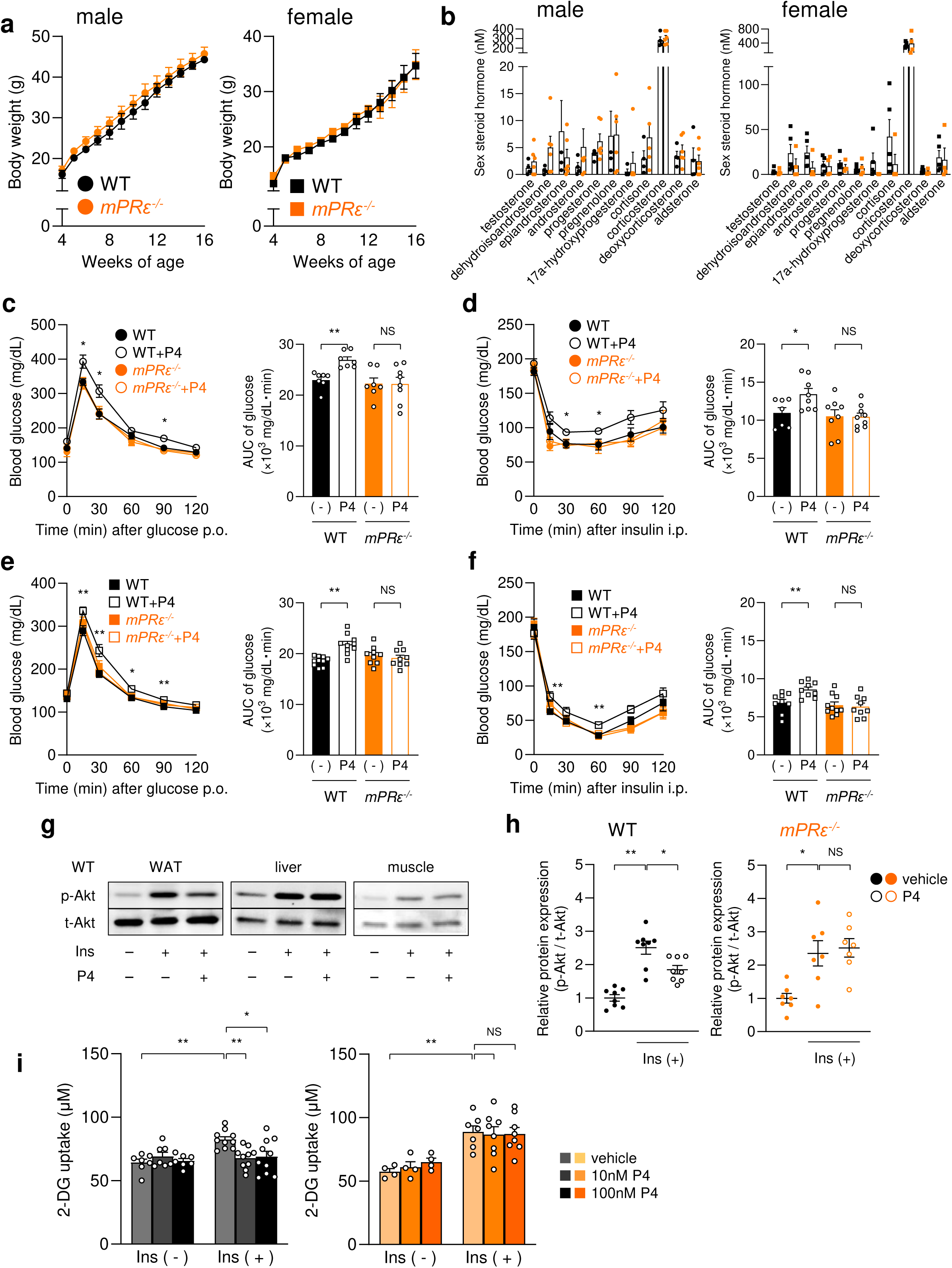
Progesterone enhances insulin resistance via adipose mPRε. (a) Changes in body weight of *mPRε^-/-^*mice under high-fat diet feeding (n = 4–6). (b) Plasma steroid levels at 16 weeks of age (n = 4–6). (c, e) Oral glucose tolerance test (OGTT) in male (c) or female (e) WT and *mPRε^-/-^* mice was performed at 15 min post-progesterone subcutaneous injection. (n = 7–10, *P < 0.05, vs. WT: progesterone). (d, f) Insulin tolerance test (ITT) in male (d) or female (f) WT and *mPRε^-/-^* mice was performed at 15 min post-progesterone subcutaneous injection (n = 7–10, *P < 0.05 vs WT: progesterone). (g) Insulin-stimulated Akt phosphorylation at Ser473 in the WATs, muscles, and liver of wild-type mice after 5 h of fasting. (h) Inhibitory effects of progesterone on insulin signaling. After pretreatment with progesterone for 30 min, a bolus of insulin (0.5 U/kg, i.p.) with or without progesterone (5 mg/kg, s.c.) was administered. Akt phosphorylation of Ser473 in WAT of wild-type mice after 5 h of fasting (n = 7–9). (i) Effect of progesterone on glucose uptake in mouse embryonic fibroblast (MEF)-derived adipocytes from WT or *mPRε^-/-^* mice. Glucose uptake was determined by measuring 2-deoxyglucose uptake using an enzymatic photometric assay (n = 4–10. *P < 0.05 vs vehicle); P4: progesterone. **P < 0.01, *P < 0.05 (Student’s t-test). The results are presented as the mean ± standard error of the mean (SE). N.S.: not significant.

To clarify whether mPRε in WATs is responsible for the observed effects of progesterone, we analyzed insulin signaling using insulin-induced Akt phosphorylation and MEF-derived adipocytes. Progesterone administration (s.c.) suppressed insulin-induced Akt phosphorylation in the WAT, but not liver or muscle tissues, of wild-type mice (Fig. 2g); however, this effect was not observed in the WAT of mPRε-deficient mice (Fig. 2h). Adipocyte differentiation from MEF, induced by a methylisobutylxanthine, dexamethasone, pioglitazone, and insulin cocktail^31^, did not differ significantly between wild-type and mPRε-deficient mice (Extended Data Fig. 3c); the influence of progesterone on adipogenesis was also comparable between wild-type and mPRε-deficient mice (Extended Data Fig. 3d). Although insulin-induced glucose uptake in MEF-induced adipocytes derived from wild-type mice was suppressed by progesterone, these effects were abolished in MEF-induced adipocytes derived from mPRε-deficient mice (Fig. 2i). These findings suggest that the activation of mPRε by progesterone in WAT promotes insulin resistance.

### Maternal insulin resistance via mPRε in pregnancy provides nutrients to embryos

Maternal insulin resistance increases in pregnancy^32^. As maternal progesterone levels also increase markedly in pregnancy, we investigated the insulin sensitivity of mPRε-deficient mice during pregnancy to clarify the relationship between progesterone and maternal insulin resistance. The progesterone levels of pregnant females (∼300 nM) were markedly higher than those of non-pregnant females (Fig. 3a). As expected, according to the glucose tolerance test, mPRε-deficient female mice on gestation day 16.5 (GD16.5) exhibited better insulin resistance than that of wild-type GD16.5 mice (Fig. 3b). However, their plasma progesterone and insulin levels were comparable (Extended Data Fig. 4a and b). Although *mPRε* mRNA expression was comparable between non-pregnant females and GD16.5 mice (Extended Data Fig. 4c), the blood glucose of GD16.5 mPRε-deficient mice was significantly lower than that of GD16.5 wild-type mice (Fig. 3c). Similarly, the blood glucose of embryos in mPRε-deficient mice was also significantly lower than that of wild-type mice (Fig. 3c). Conversely, the insulin sensitivity of non-pregnant mPRε-deficient mice was similar to that of non-pregnant wild-type mice (Fig. 3d).

**Fig. 3.**
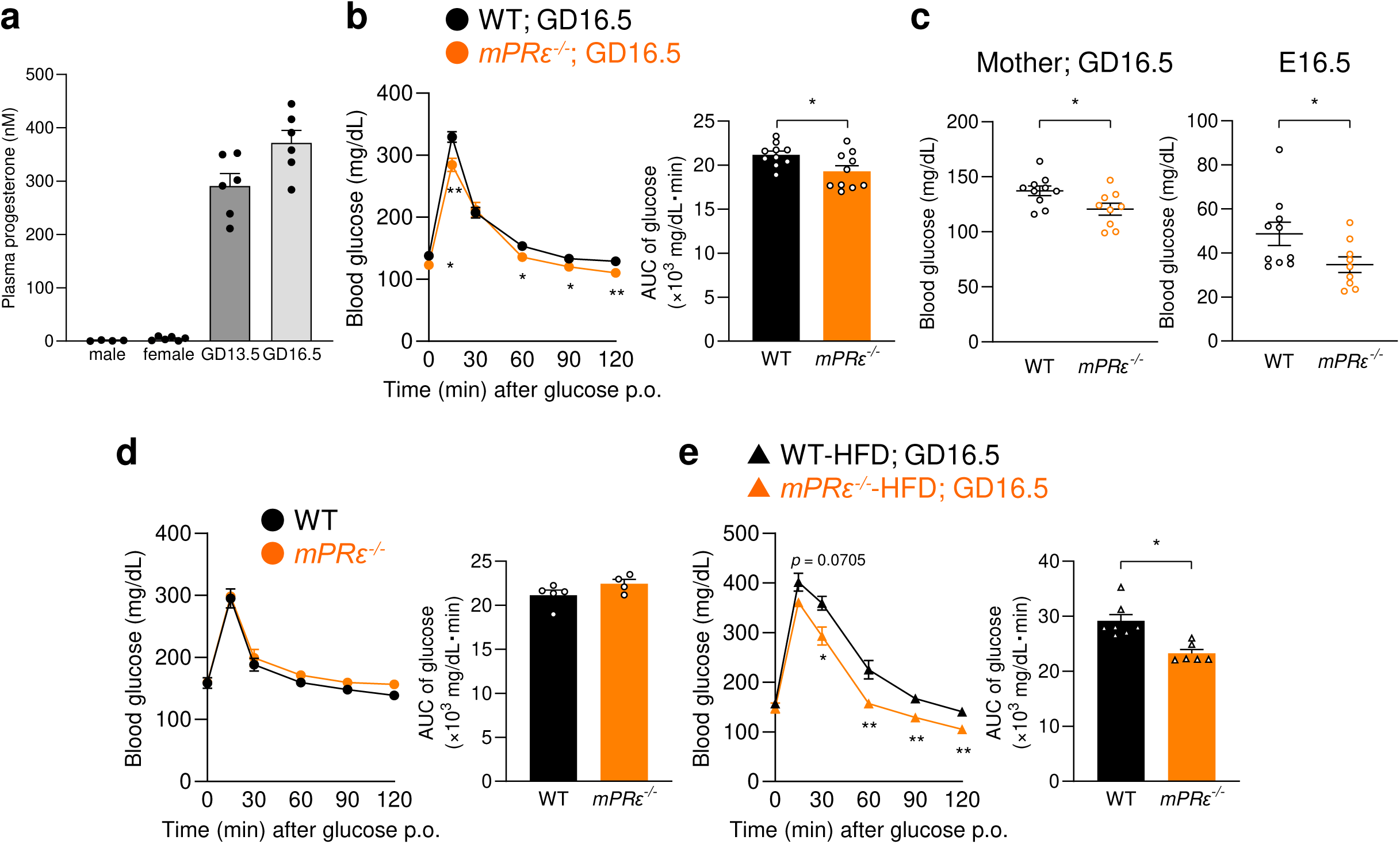
Maternal progesterone-adipose mPRε in pregnancy control blood glucose of embryos. (a) Plasma progesterone levels in mice (males, non-pregnant, GD13.5, and GD16.5 pregnant mice) (n = 4–6). (b) oral glucose tolerance test (OGTT) in GD16.5 wild-type and *mPRε*^-/-^ mothers (n = 10, *P <0.05, **P <0.01 vs WT). (c) Blood glucose levels in GD16.5 WT and *mPRε*^-/-^ mothers (left) and E16.5 embryos (right) (n = 9–10, **P <0.01 vs WT). (d) OGTT in non-pregnant mice n = 4–5, *P <0.05, **P <0.01 vs WT) (e) OGTT in HFD-induced GDM model GD16.5 wild-type and *mPRε*^-/-^ mothers (n = 6–7). **P < 0.01, *P < 0.05 (Mann–Whitney U test). Results are presented as the mean ± standard error of the mean (SE).

Maintaining adequate blood glucose levels in gestational diabetes mellitus reduces morbidity for both mother and baby^6^. We analyzed insulin resistance in a mouse model of HFD-induced gestational diabetes mellitus and found that insulin resistance in GD16.5 wild-type mice was markedly improved compared to that in GD16.5 mPRε-deficient mice (Fig. 3e). Thus, mPRε-induced maternal insulin resistance during pregnancy enhances glucose availability for embryos.

### mPRε deficiency in pregnancy causes postnatal growth abnormalities

We investigated the influence of maternal mPRε on offspring growth (Fig. 4a). On postnatal day 28, the body weights of both mPRε-deficient and heterozygous offspring from mPRε-deficient mothers were significantly lower than those of wild-type and heterozygous offspring from wild-type mothers (Fig. 4b). Moreover, mPRε-deficient mice from two mPRε-deficient parents and wild-type mice from two wild-type parents were fed a high-fat diet (HFD) during growth. Unlike mPRε-deficient mice from a heterozygous mother (Fig. 2a), the body weights of mPRε-deficient mice from a mPRε-deficient mother were significantly lower than those of wild-type mice from a wild-type mother for both males and females during growth (Fig. 4c). Similarly, the liver and WAT weights of mPRε-deficient male mice and the WAT weights of mPRε-deficient female mice from mPRε-deficient mothers at 16 weeks of age were also significantly lower than those of wild-type mice from wild-type mothers (Fig. 4d). The levels of plasma triglycerides, non-esterified fatty acids, and total cholesterol in mPRε-deficient male mice (Extended Data Fig. 5a) from mPRε-deficient mothers at 16 weeks of age, and the levels of plasma non-esterified fatty acids and total cholesterol in mPRε-deficient female mice (Extended Data Fig. 5c) from mPRε-deficient mothers at 16 weeks of age were also significantly lower than those of wild-type mice from wild-type mothers. Notably, the plasma insulin levels of mice from mPRε-deficient mothers at 16 weeks of age were significantly lower (in males; Extended Data Fig. 5b) and higher (in females; Extended Data Fig. 5e) than those of wild-type mice from wild-type mothers. Furthermore, plasma progesterone levels were less than 10 nM in both males and females (Extended Data Fig. 5c, f). Thus, the offspring of mPRε-deficient mothers exhibit metabolic dysfunction related to leanness.

**Fig. 4.**
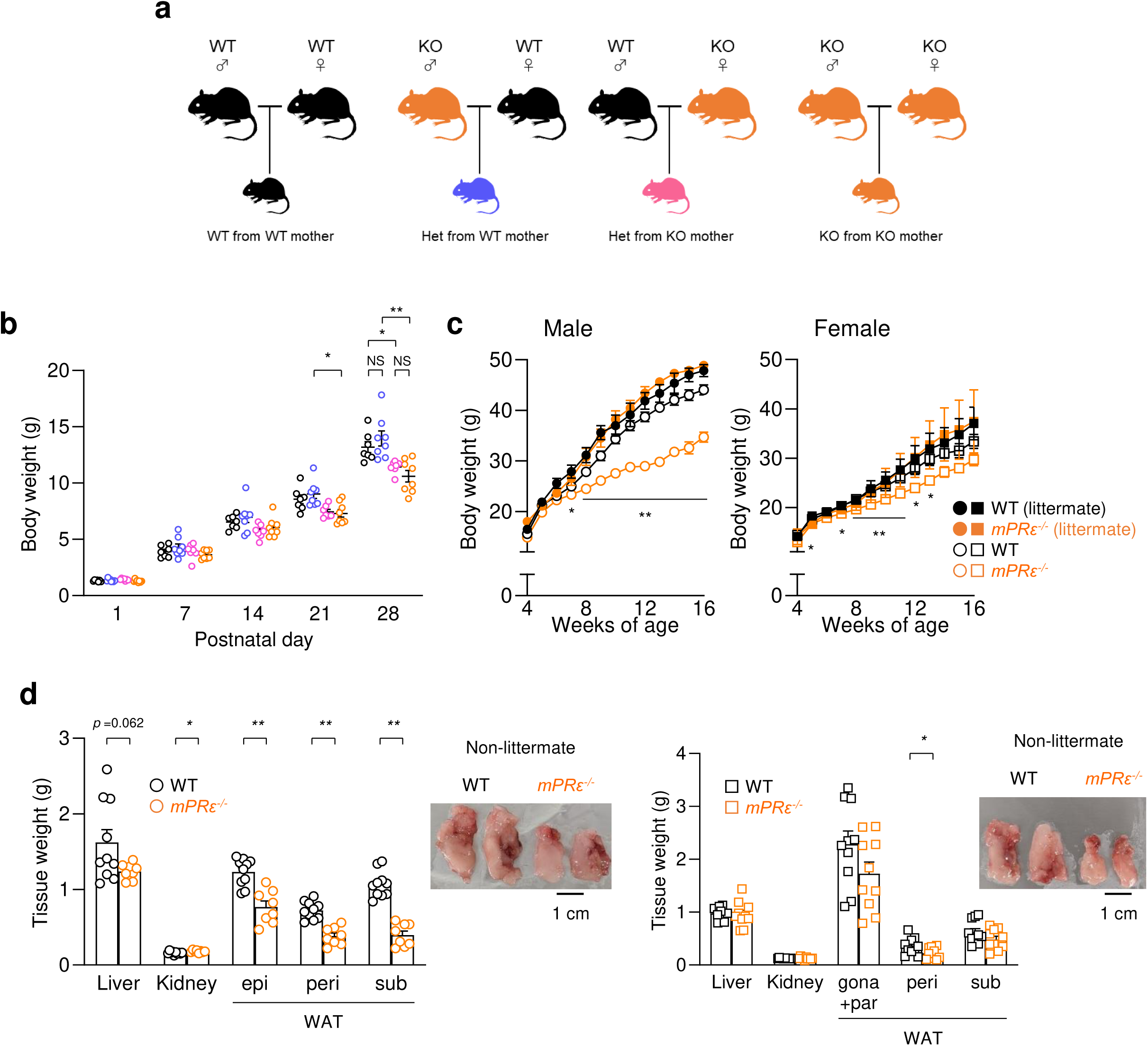
Offspring from mPRε deficient mother shows metabolic dysfunctions. (a) Experimental scheme. (b) Body weight change during the growth of pups born from homozygous crosses (n = 9–10, *P <0.05, **P <0.01 vs WT from WT). Body weight change during growth pups born from heterozygous crosses (n = 4–5, **P <0.05, *P <0.01 vs Hetero from wild-type) (c) Changes in body weight of *mPRε^-/-^* mice under high-fat diet feeding in males (left) or females (right) (n = 4–10). (d) The tissue weights of *mPRε^-/-^* mice at 16 weeks of age under high-fat diet feeding in males (left) or females (right) (n = 8–10). **P < 0.01, *P < 0.05 (Mann–Whitney U test). Results are presented as the mean ± standard error of the mean (SE). N.S.: not significant.

### Adipose mPRε exerts insulin resistance via prostaglandin production

We investigated intracellular signaling related to adipose mPRε-mediated insulin resistance. RNA sequencing and KEGG enrichment analyses were performed to compare the WAT of GD16.5 and non-pregnant wild-type female mice. The results revealed a relationship between the prostaglandin synthesis pathway, chronic inflammation, and insulin resistance-related pathways, which were not observed in the WAT of mPRε-deficient mice (Fig. 5a–d) or in the livers of wild-type mice (Extended Data Fig. 6a, b). Similarly, lipid metabolome profiling of WAT in GD16.5 and non-pregnant female mice revealed significant alterations in lipid mediators and polyunsaturated fatty acids, which are related to inflammation and insulin resistance (Fig. 5e). Moreover, quantitative analysis revealed that prostaglandin E_2_ (PGE_2_) and arachidonic acid levels in the WAT of mPRε-deficient GD16.5 mothers were significantly lower than those in wild-type GD16.5 mothers (Fig. 5f). Prostaglandins via the arachidonic acid cascade are considered lipid mediators of the inflammatory response that promote insulin resistance in WAT^33,34^.

**Fig. 5.**
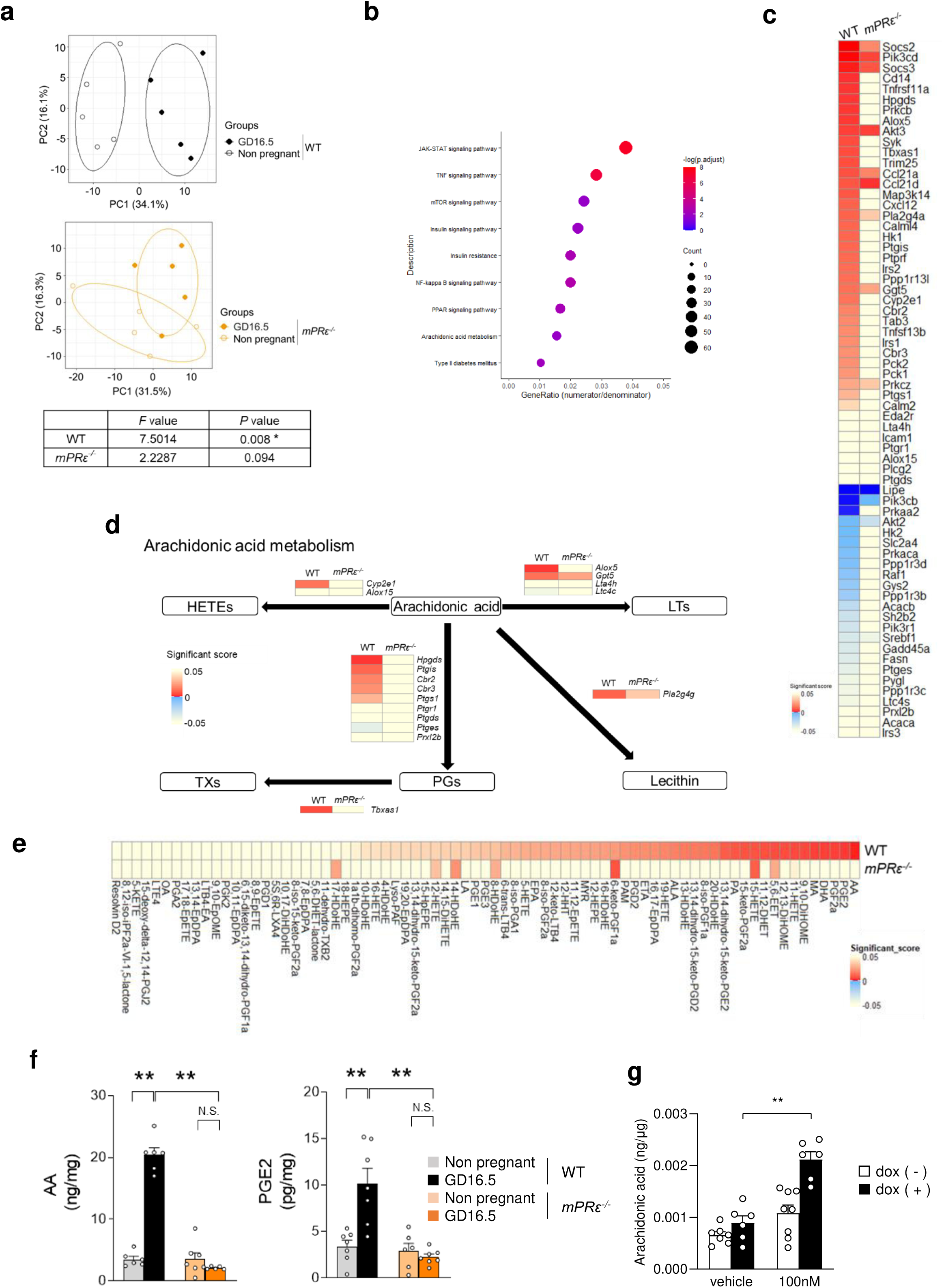
Production of prostaglandins increased in pregnancy via mPRε of WAT. (a) The beta diversity as shown via the principal component analysis (PCA) based on genes from KEGG (ID mmu00590; mmu04064; mmu04910) in the WATs of WT and *mPRε^-/-^* mice with non-pregnant or GD16.5 (n = 5). (b) KEGG enrichment analysis related to molecular function in WATs of GD16.5 *mPRε^-/-^* mother. P-values were adjusted based on the false discovery rate (FDR). (c) Heat map of prostaglandin synthesis pathway, chronic inflammation, insulin resistance-related gene profiles of the WATs of WT and *mPRε^-/-^* mice with non-pregnant or GD16.5 (n = 5). Among non-pregnant vs GD16.5 mother, 66 identified genes demonstrated significant differences in abundance (absolute log2 fold change > 0.5, p < 0.05: red open square). (d) KEGG pathway analysis related to arachidonic acid metabolism in WATs of GD16.5 *mPRε^-/-^*mothers. P-values were adjusted based on the false discovery rate (FDR). (e) Comprehensive analysis of lipid mediators in WATs of GD16.5 *mPRε^-/-^*mothers. Heat map of the top 40% relative lipid metabolites profiles of the WATs of WT and *mPRε^-/-^* mice with non-pregnant or GD16.5 (n = 5). Among non-pregnant vs GD16.5 mother, 75 identified genes demonstrated significant differences in abundance (absolute log2 fold change > 0.5, p < 0.05: red open square). (f) Arachidonic acid (left) and PGE_2_ (right) were quantified in the WATs of non-pregnant mice or GD16.5 mothers. (n = 6–7 per group for arachidonic acid; n = 6–7 per group for PGE_2_). (g) Arachidonic acid levels in response to progesterone (100 nM) in Flp-In mPRε T-REx HEK293 cells treated with or without doxycycline (n = 6–8). Dox; doxycycline. **P < 0.01, *P < 0.05 (Mann–Whitney U test). The results are presented as the mean ± standard error of the mean (SE). N.S.: not significant.

The mPR family subtype mPRβ influences release of fatty acids from membrane phospholipids^27^. Therefore, we compared the profiles of membrane phospholipids in the WAT of mice with and without pregnancy. Phosphatidylserine in the membrane phospholipids of WAT of pregnant wild-type mice exhibited decreased high-carbon-chain fatty acids and increased low-carbon-chain fatty acids, which was not observed in mPRε-deficient mice (Extended Data Fig. 7a, 8). Additionally, phospholipase A_2_ (PLA_2_) activity in the WAT of mPRε-deficient GD16.5 mothers was significantly lower than that in wild-type GD16.5 mothers (Extended Data Fig. 7b).

We further examined the effects of progesterone on mPRε-mediated release of fatty acids from membrane phospholipids in the heterologous expression system. Similar to reports on other mPR family subtypes^27^, progesterone promoted the phosphorylation of ERK but not other GPCR signaling pathways such as intracellular cAMP concentration in HEK293 cells expressing mouse mPRε (Extended Data Fig. 9a–c). Progesterone significantly enhanced PLA_2_ activity in mPRε-overexpressing HEK293 cells, whereas these effects were not observed in doxycycline-uninduced control [Dox (−)], non-mPRε-expressing HEK293 cells (Extended Data Fig. 7c). Additionally, progesterone stimulation increased arachidonic acid levels in mPRε-overexpressing HEK293 cells, whereas these effects were not observed in Dox (−) control HEK293 cells (Fig. 5g). Thus, mPRε activation by progesterone in WAT promotes PGE_2_ synthesis by releasing arachidonate from phospholipids, thereby increasing insulin resistance.

## Discussion

Although the physiological functions of the membrane progesterone receptor mPRε indicate an influence on energy metabolism, the relationship between mPRε and progesterone as a ligand for mPRε remains unclear. In this study, we demonstrate that mPRε, which was significantly expressed in WAT of the mPR family, enhanced adipose insulin resistance via prostaglandin production through the release of arachidonate from membrane phospholipids by progesterone-stimulated PLA_2_ activation. This increase in maternal insulin resistance caused by the interaction between progesterone and adipose mPRε during pregnancy controlled the provision of glucose to embryos and growth after birth.

We suggest that the role of progesterone in adipocytes depends on mPRε because mPRε was the only member of the mPR family that was abundantly expressed in the WAT of both males and females. Although the nuclear progesterone receptor was also expressed in WAT, expression was localized to only the stromal vascular fraction in females. Moreover, our finding that progesterone administration increased insulin resistance via mPRε in male mice, as well as female mice, implies that mPRε may have important functions regardless of gender. Progesterone, but not estrogen, is a precursor of all steroids in the steroid biosynthetic pathway; therefore, under some physiological conditions, progesterone may locally, partially, and temporarily reach high concentrations, even in males. Alternatively, the fact that mPRα binds to various steroids, albeit with lower affinity than with which it binds to progesterone^34^, suggests that mPRε may be activated by steroid metabolites other than progesterone.

mPRε affects energy metabolism via the liver and pancreas^28,29^. Therefore, during pregnancy, mPRε present in other tissues besides WAT, such as the liver and pancreas, may also influence maternal insulin sensitivity. However, our results showed that progesterone administration suppressed Akt phosphorylation related to the insulin signaling pathway in WAT but not in liver or muscle tissue and did not affect plasma insulin levels. Moreover, in pregnancy, progesterone administration did not affect maternal plasma insulin levels, and RNA seq showed that the expressions of lipid, glucose, and energy metabolism-related genes in the liver were comparable between wild-type and mPRε-deficient mothers. However, progesterone in pregnancy may induce other functions of mPRε besides blood glucose control between the mother and embryo and may also induce functions of other receptors. Further studies are required to clarify progesterone-induced interactions between mother and embryo.

The offspring of mPRε-deficient mothers exhibited phenotypes of leanness under HFD feeding in this study. Low birth weights caused by a decrease in glucose provision to embryos through increased insulin sensitivity may be the reason for this phenotype. According to previous research, low birth weight leads to metabolic dysfunctions such as obesity or other symptoms related to leanness during growth^3–12^. However, the mechanism remains unclear. Our results suggest that variations in maternal progesterone levels during pregnancy may modulate the optimal concentration of embryonic blood glucose, thereby leading to metabolic phenotypes in offspring. Nevertheless, further studies are required to clarify the impact of longitudinal maternal glycemic control during pregnancy on the development of embryos and offspring growth after birth.

Our results demonstrated that mPRε activation by progesterone promotes intracellular PGE_2_ and arachidonate production synthesis. This progesterone-mPRε-mediated response may be caused by the release of arachidonate from phospholipids via PLA_2_ activity; particularly, it reported that mPRβ is associated with PLA_2_ activity^27^. Our lipid metabolome indicated that other fatty acids as well as arachidonate also increased in WAT along with pregnancy. These results suggest that not only glucose but also NEFAs may be efficiently supplied to embryos via adipose mPRε activation. However, further studies need to clarify the progesterone-mPRε-mediated intracellular signaling mechanism because the accurate interaction between mPRβ and PLA_2_ activity is until unclear.

Additionally, our studies predominantly focused on murine models; hence, their translational relevance to human physiology remains to be clarified.

In conclusion, we showed that adipose mPRε activated by high concentrations of maternal progesterone during pregnancy increased maternal insulin resistance, ensuring the efficient provision of glucose to embryos. Dysfunctions of maternal blood glucose control in pregnancy can lead to low birth weight or macrosomia, which subsequently increases the risk for metabolic diseases after birth. Therefore, our results have implications for the development of mPRε-targeting drugs for the selective treatment of such metabolic diseases, as well as for suppressing side effects without targeting progesterone.

## Acknowledgments

This work was supported by research grants from the AMED (JP23gm1510011 to I.K), JST-MOONSHOOT (JPMJMS2023 to I.K. and J.A.).

## Author contributions

K.W. performed the experiments and wrote the paper; M.Y. performed the experiments and wrote the paper; J.M. performed the experiments and interpreted the data; R.O.K. performed the experiments and interpreted the data; Y.M. performed the experiments and interpreted the data; D.S. performed the experiments; Y.M. performed the experiments; N.K. performed the experiments; S.I. performed interpreted the data; F.O. performed the experiments; K.N. interpreted the data; J.A. interpreted the data; Y.S. performed the experiments; H.O. interpreted the data; E.K. interpreted the data; I.K. supervised the project, interpreted the data, and wrote the paper; I.K. had primary responsibility for the final content. All authors read and approved the final version of the manuscript.

## Data availability

The source data presented in RNA-seq analysis have been deposited in the DNA Data Bank of Japan (DDBJ) under the accession nos. E-GEAD-852, E-GEAD-853, and E-GEAD-854. The source data presented in Figures 1–5 and Extended Data Figures 1–9 have been deposited into the Dryad repository (https://doi.org/10.5061/dryad.280gb5mzf).

## Methods

### Animals

Male C57BL/6 mice were obtained from Japan SLC (RRID: IMSR_JAX:000664, Shizuoka, Japan), while *mPRε^−/−^* mice with C57BL/6N background were generated. The mice were kept in a conventional animal facility at 24 °C with a 12 h light/dark cycle and were acclimated to the CLEA Rodent Diet (CE-2; CLEA Japan, Inc., Tokyo, Japan) for 1 week before starting the treatments. For the high fat diet (HFD) feeding study, 4-week-old mice were fed either NC or HFD with 60% kcal fat (D12492, Research Diets Inc., New Brunswick, NJ, USA) for 12 weeks. Body weight was recorded weekly throughout the experiment. At the end of the study, all the mice were euthanized under deep isoflurane anesthesia. All procedures involving animals were conducted in compliance with the guidelines of the Committee on the Ethics of Animal Experiments of the Kyoto University Animal Experimentation Committee (Lif-K23012), ensuring that all efforts were made to minimize animal distress.

### RNA extraction and real-time quantitative RT-PCR

Total RNA was extracted using an RNeasy Mini Kit (Qiagen, Hilden, Germany) and RNAiso Plus reagent (TAKARA). Complementary DNA (cDNA) was synthesized from the RNA templates using Moloney murine leukemia virus reverse transcriptase (Invitrogen, Carlsbad, Cam USA). Quantitative reverse transcriptase PCR (qRT-PCR) was conducted using SYBR Premix Ex Taq II (TAKARA) on a StepOnePlus real-time PCR system (Applied Biosystems, Foster City, CA, USA), following previously described protocols^31^. The PCR conditions were as follows: initial denaturation at 95°C for 30 s, followed by 40 cycles of 95 °C for 5 s, 58 °C for 30 s and 72 °C for 1 min. The dissociation stage was carried out at 95 °C for 15 s, then 60 °C for 1 min, and a final step at 95 °C for 15 s. Primer sequences were as follows: mPRε, 5′-CACTTCATCCCGCTGCTGCT-3′ (forward) and 5 ′ - GCGGCTCTTACAGCAAGCCA -3 ′ (reverse); mPRα, 5 ′-CGGCATGGCGATGGCAGTA-3 ′ (forward) and 5 ′ -CTGCACCTTGTCATGCCAGG-3 ′ (reverse); mPRβ, 5 ′ -CACCGCTGTGTCATGACGCT-3 ′ (forward) and 5 ′-GCCTGGTCGGAGCTATAGA-3′ (reverse); mPRγ, 5′-TGACAGCTACTCGTGGCCGA-3′ (forward) and 5 ′ -GCCCATGTGCTTCTGGTGGT-3 ′ (reverse); mPRδ, 5 ′ -TACTGCCCGCCTGCCTCTAT-3 ′ (forward) and 5 ′-TGAGAGCAGAGCGCAGAGGA-3 ′ (reverse); PGR, 5 ′ -CGACGTGGAGGGAGCTTTCT-3 ′ (forward) and 5 ′-CCTGGGTGGTGACAGTCCTT-3 ′ (reverse); and 18S, 5 ′-ACGCTGAGCCAGTCAGTGTA-3 ′ (forward) and 5 ′ -CTTAGAGGGACAAGTGGCG-3 ′ (reverse).

### Biochemical analyses

Plasma levels of non-esterified fatty acid (NEFA) (LabAssay™ NEFA; FUJIFILM Wako Pure Chemical Corporation, Osaka, Japan), triglyceride (LabAssay™ Triglyceride; FUJIFILM Wako Pure Chemical Corporation), and total cholesterol (LabAssay™ Cholesterol; FUJIFILM Wako Pure Chemical Corporation) in mice were measured following the manufacturer’s protocols. Blood glucose levels were determined using a handheld glucometer (OneTouch Ultra; LifeScan, Milpitas, CA, USA). Plasma insulin levels were assessed using an insulin ELISA kit [insulin enzyme-linked immunosorbent assay (ELISA) kit (RTU); Shibayagi, Gunma, Japan], per the manufacturer’s instructions.

### Steroids measurement

For steroid measurements, plasma samples were mixed with methanol containing an internal standard, followed by ethyl acetate for lipid extraction. The mixture was centrifuged at 8,000 g at 4 °C for 15 min, and the supernatant containing the steroids was collected and dried. Subsequently, the dried sample was redissolved in methoxyamine hydrochloride (20 mg/mL in pyridine) and incubated (60 °C, 45 min) for derivatization. The dried samples were resuspended in acetonitrile and analyzed by liquid chromatography-tandem mass spectrometry (LC-MS/MS). This analysis was performed using an ultra-performance liquid chromatography (UPLC) system (Waters, Milford, MA, USA) equipped with an Acquity UPLC system coupled to a Waters Xevo TQD mass spectrometer (Waters). Separation was achieved using a acetonitril gradient in a 0.1% formic acid aqueous solution on an ACQUITY UPLC BEH C18 column (2.1 × 150 mm, 1.7 μm; Waters).

### Insulin sensitivity analysis

To evaluate glucose tolerance, mice were given an oral gavage of glucose (1.5 g/kg body weight) after a 16-h fast. For the insulin tolerance test, mice were fasted for 4 h and then intraperitoneally injected with insulin (0.5 U/kg; Sigma-Aldrich, St. Louis, MO, USA). Blood glucose levels were measured before injection and at 15, 30, 60, 90, and 120 min post-injection. To assess the biochemical responses to insulin stimulation, mice were injected intraperitoneally with insulin (0.5 U/kg). After 15 min, the liver, skeletal muscle, and WATs were dissected and immediately frozen in liquid nitrogen. Immunoblotting was performed as previously described^31^.

### Western blotting

Tissues were homogenized in 0.1 M sodium phosphate buffer (pH 7.4) and centrifuged at 14,000 g for 30 min at 4 °C. Tissue lysates were prepared in a TNE buffer containing 10 mM Tris-HCl (pH 7.4), 150 mM NaCl, 1 mM EDTA, 1% Nonidet P-40, 50 mM NaF, 2 mM Na_3_VO_4_, 10 μg/mL aprotinin, and a 1% phosphatase inhibitor cocktail (Nacalai Tesque, Kyoto, Japan). Proteins from the lysates were separated using sodium dodecyl sulfate-polyacrylamide gel electrophoresis and transferred to nitrocellulose membranes. Proteins, such as Akt, ERK, and their activated forms, were detected by western blotting using specific antibodies. The primary antibodies used were Akt (1:1000) (Cell Signaling Technology, Danvers, MA, USA), phosphorylated Akt (1:1000) (Cell Signaling Technology), ERK (1:1000) (Cell Signaling Technology), and phosphorylated ERK (1:1000) (Cell Signaling Technology). The secondary antibody was used a horseradish peroxidase-conjugated donkey anti-rabbit antibody (1:2000) (GE Healthcare). Immunoreactive bands were visualized using an enhanced chemiluminescence detection system as previously described^31^. The ImageJ software (National Institutes of Health) was used to quantify the integrated density of each band.

### Plasma glucose measurement

To quantify glucose in the embryo plasma, acetone was added to the plasma and mixed using a vortex mixer. The mixture was centrifuged at 10,000g for 5 min at 4 °C, and the supernatant was collected for LC-MS/MS analysis. Glucose was analyzed using an Acquity UPLC system coupled to a Waters Xevo TQD mass spectrometry (Waters) and separated on an ACQUITY UPLC BEH Amide column (2.1 × 150 mm, 1.7 μm; Waters) using a concentration gradient of water, acetonitrile, acetone, and 4% ammonium hydroxide.

### Adipocytes culture

MEF-derived adipocytes were cultured at 37 °C in α-MEM supplemented with 1% penicillin-streptomycin solution (Gibco) and 10% fetal bovine serum (FBS). Then, 2 d after reaching confluence, the medium was replaced with α-MEM containing 10% FBS and inducers (0.25 μM dexamethasone, 10 μg/mL insulin, and 0.5 mM 3-isobutyl-1-methylxanthine, IBMX) along with pioglitazone (10 μM) for 2 d. Subsequently, the medium was switched to DMEM supplemented with 10% FBS, 10 μg/mL insulin, and 10 μM pioglitazone for another 12 d. After 12 d, the fully differentiated adipocytes were ready for experimentation^31^. For the glucose uptake assay, MEF-derived adipocytes were incubated in a serum-free medium for 6 h, followed by the addition of insulin (3 μg/mL) and incubation at 37 °C for 20 min. Subsequently, 0.1 mM 2-deoxyglucose (Sigma) and progesterone were added, and the cells were incubated at 37 °C for an additional 20 min. The adipocytes were then washed thrice with ice-cold PBS containing phloretin and collected in 1% NP-40/10 mM Tris-HCl. The cell lysates were heat-treated at 80 °C for 15 min, centrifuged at 15,000 g for 20 min at 4 °C, and the supernatants were collected. The collected supernatants were analyzed using a 2-deoxyglucose (2DG) Uptake Measurement kit (Cosmo Bio). For Oil Red O staining, the cells were fixed with 4% PFA for 10 min and washed thrice with PBS. Fixed cells were stained with Oil Red O for 10 min, washed with MQ, and extracted with isopropanol. The absorbance was measured for quantitative analysis.

### RNA sequencing

RNA was extracted from the WAT and liver of non-pregnant and pregnant mice using RNAiso Plus reagent (TAKARA) and the RNeasy Mini Kit (Qiagen). RNA integrity, quality, and concentration were measured using an Agilent 2100 Bioanalyzer system with an RNA 6000 Nano Kit (Agilent Technologies). Subsequently, RNA sequencing libraries were prepared using the NEBNext® Ultra™ II Directional RNA Library Prep Kit (Illumina) and NEBNext Multiplex Oligos for Illumina (Dual Index Primers Set 1) and sequenced on an Illumina NovaSeq 6000. Each sample yielded approximately four gigabases of paired-end reads with a length of 150 bp. The obtained RNA sequencing data were processed using trimmomatic-0.39 to eliminate adapters and low-quality reads^36^. The quality of the trimmed sequences was evaluated using FastQC (version 0.11.8.-2)^37^. Alignment of the reads to the mouse reference genome (NCBI GRCm39) was performed using the STAR software (version 2.7.10a)^38^. The raw read counts were subjected to relative log expression normalization to identify differentially expressed genes (DEGs) across all comparisons. Fold changes were calculated using RSEM (version 1.3.3) and edgeR^39^. DEGs were determined based on two criteria: a false discovery rate (FDR)-adjusted p-value < 0.05 (Benjamini-Hochberg procedure) and an absolute log2 fold change > 0.5. Gene Set Enrichment Analysis was performed using the Kyoto Encyclopedia of Genes and Genomes (KEGG) database (http://www.genome.jp/kegg/).

### Fatty acids measurement

To quantify individual FAs, samples (approximately 50 mg adipose tissue or 1.2 × 10^6^ mPRε-overexpressing HEK293 cells) were homogenized in methanol (1 mL) with an internal control (C17:1). Chloroform (2 mL) and 0.5 M potassium chloride (0.75 mL) were then added to extract lipids. The collected lipid layers were dried and the samples were resuspended in chloroform:methanol (1:3, v/v) for LC-MS/MS analysis. FAs were analyzed using an Acquity UPLC system coupled to a Waters Xevo TQD mass spectrometry (Waters) and separated on an ACQUITY UPLC BEH C18 column (2.1 × 150 mm, 1.7 μm; Waters) using an acetonitrile gradient in 10mM ammonium formate aqueous solution. The flow rate was 0.4 mL/min, and the column temperature was maintained at 50 °C. MS detection was performed in the negative ionization mode, with the source capillary voltage set to 3000 V. The desolvation and source temperatures were set at 500 °C and 150 °C, respectively. Individually optimized multiple reaction monitoring parameters were determined for the target compounds using the standards.

### Comprehensive analysis of lipid mediators

To quantify prostaglandins in tissues, solid-phase extraction was performed using Oasis HLB cartridges (1 mg; Waters, Milford, MA, USA), according to the method described by Lee et al^40^. Briefly, the samples were homogenized in methanol and the supernatants were diluted with water to achieve a final methanol concentration of approximately 7%. The diluted samples were applied to pretreated cartridges and sequentially washed with 0.1% formic acid, 15% ethanol, and hexane. The samples were eluted with 200 μL of methanol and dried. The dried samples were subsequently reconstituted in 20 μL of methanol and subjected to LC-MS/MS analysis. A Shimadzu LC/MS/MS Method Package for Lipid Mediators ver. 2 (Shimadzu) was used to analyze various lipid mediators. The analysis was performed on a Shimadzu LCMS-8060NX Triple Quadrupole LC-MS/MS system and separated on a Kinetex C8 column (2.1 × 150 mm, 2.6 μm; Phenomenex).

### Phospholipids measurement

Total lipids in WAT samples were extracted using the method described by Bligh and Dyer^41^. The extracted lipids were dried up with a centrifugal evaporator, dissolved in methanol: isopropanol=1:1, and stored at −20 °C. Lipid samples were subjected to LC/ESI-MS-based lipidomic analyses using a Shimadzu Nexera UPLC system (Shimadzu) coupled with a QTRAP 4500 hybrid triple quadrupole linear ion trap mass spectrometer (SCIEX). Chromatographic separation was performed on a SeQuant ZIC-HILIC PEEK coated column (250 mm × 2.1 mm, 1.8 µm; Millipore) maintained at 50 °C using mobile phase A (water/acetonitrile (95/5, v/v) containing 10 mM ammonium acetate) and mobile phase B (water/acetonitrile (50/50, v/v) containing 20 mM ammonium acetate) in a gradient program (0–22 min: 0% B→40% B; 22–25 min: 40% B→40% B; 25–30 min: 0% B) with a flow rate of 0.3 mL/min. Instrument parameters were as follows: curtain gas, 30 psi; collision gas, 7 arb. unit; ionspray voltage, −4500 V; temperature, 700 °C; ion source gas 1, 30 psi; ion source gas 2, 70 psi. Phospholipid species were detected by multiple reaction monitoring (MRM) as previously described^42^.

### PLA_2_ activity in WAT

WAT samples of WT and *mPRε^−/−^* mice with GD16.5 mothers were washed with PBS solution containing 0.16 g/mL heparin (pH 7.4) to eliminate adhered red blood cells and clots. After removing PBS, samples were homogenized in 50 mM HEPES containing 1 mM EDTA buffer (pH 7.4) on ice and centrifuged at ×10,000 g for 15 min at 4 °C. The supernatants were collected, and PLA_2_ activity was measured using a phospholipase A2 calcium-dependent cytosolic assay kit (cPLA_2_ Assay Kit, Cayman Chemical, Ann Arbor, MI, USA) per the manufacturer’s instructions.

### HEK293 cells expressing mouse mPRε

Flp-In T-REx HEK293 cells were sourced from Invitrogen. To generate HEK293 cells expressing mouse mPRε, the cells were transfected with a mixture of pcDNA5/FRT/TO-E-tag-mPRε and pOG44 using Lipofectamine reagent (Invitrogen)21. Transfected cells were cultured in DMEM supplemented with 10 μg/mL blasticidin S (Funakoshi, Tokyo, Japan), 100 μg/mL hygromycin B (Gibco, Grand Island, NY, USA), and 10% FBS. The cells were maintained at 37 °C in a 5% CO_2_ atmosphere and cultured under various conditions. To assess the PLA_2_ activity, HEK293 cells were plated in 24-well plates and cultured for 24 h. Subsequently, each well was stimulated with progesterone for 24 h and collected in 50 mM HEPES buffer containing 1 mM EDTA (pH 7.4). The lysates were centrifuged at ×10,000 g for 15 min at 4°C. The supernatants were collected, and the PLA_2_ activity was measured using the cPLA_2_ Assay Kit (Cayman Chemical) following the manufacturer’s protocol. The cAMP concentration was determined by enzyme immunoassay (EIA) using a cAMP EIA kit (Cayman Chemical), following the manufacturer’s protocol. For cAMP determination, the cells were lysed in a 0.1 N HCl solution21. All assays were conducted in duplicates.

### Statistical analysis

All data are expressed as the mean ± standard error of the mean (SEM). Statistical analyses were performed using GraphPad Prism software (GraphPad Software Inc., La Jolla, CA, USA, RRID: SCR_002798). Data normality was assessed using the Shapiro–Wilk test. Depending on the normality of the data, statistical comparisons were conducted using the Student’s t-test (two-tailed), Mann–Whitney U test (two-tailed), or two-way ANOVA with the Bonferroni post hoc test. Statistical significance was defined as P < 0.05.

## Supplementary information

**Extended Data Fig. 1.**
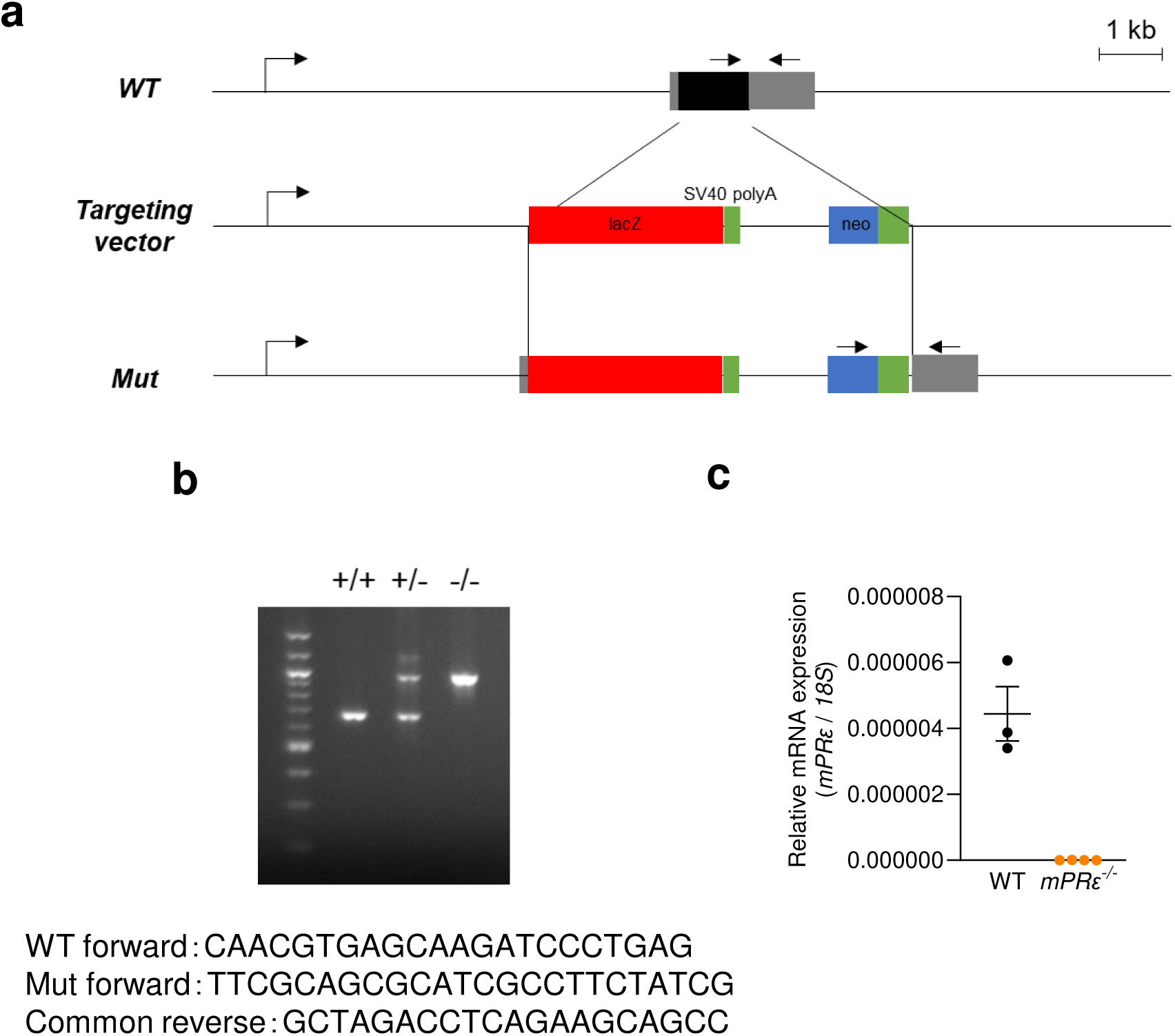
Schematic representation of the mPRε gene structure and expression. (a) A targeting vector was constructed by ligation of three fragments: the 5’ and 3’ homology recombination arms and a fragment of the LacZ-PGK-neo cassette. A 1.6-kbp fragment of mouse DNA containing the exon coding for mPRε was replaced with the LacZ-PGK-neo cassette. The linearized targeting vector was electroporated into 129/Sv embryonic stem (ES) cells. (b) Mouse genotypes were determined by PCR using three 3 primers: P1, P2, and P3 (wild-type allele: 671bp, P1/P3; mutant allele, 968bp, P1/P2). (c) Expression of *mPRε* mRNA in mouse adipose tissues of wild-type and *mPRε^-/-^* mice (n = 3–4). The expression of *18S* rRNA was used as an internal control.

**Extended Data Fig. 2.**
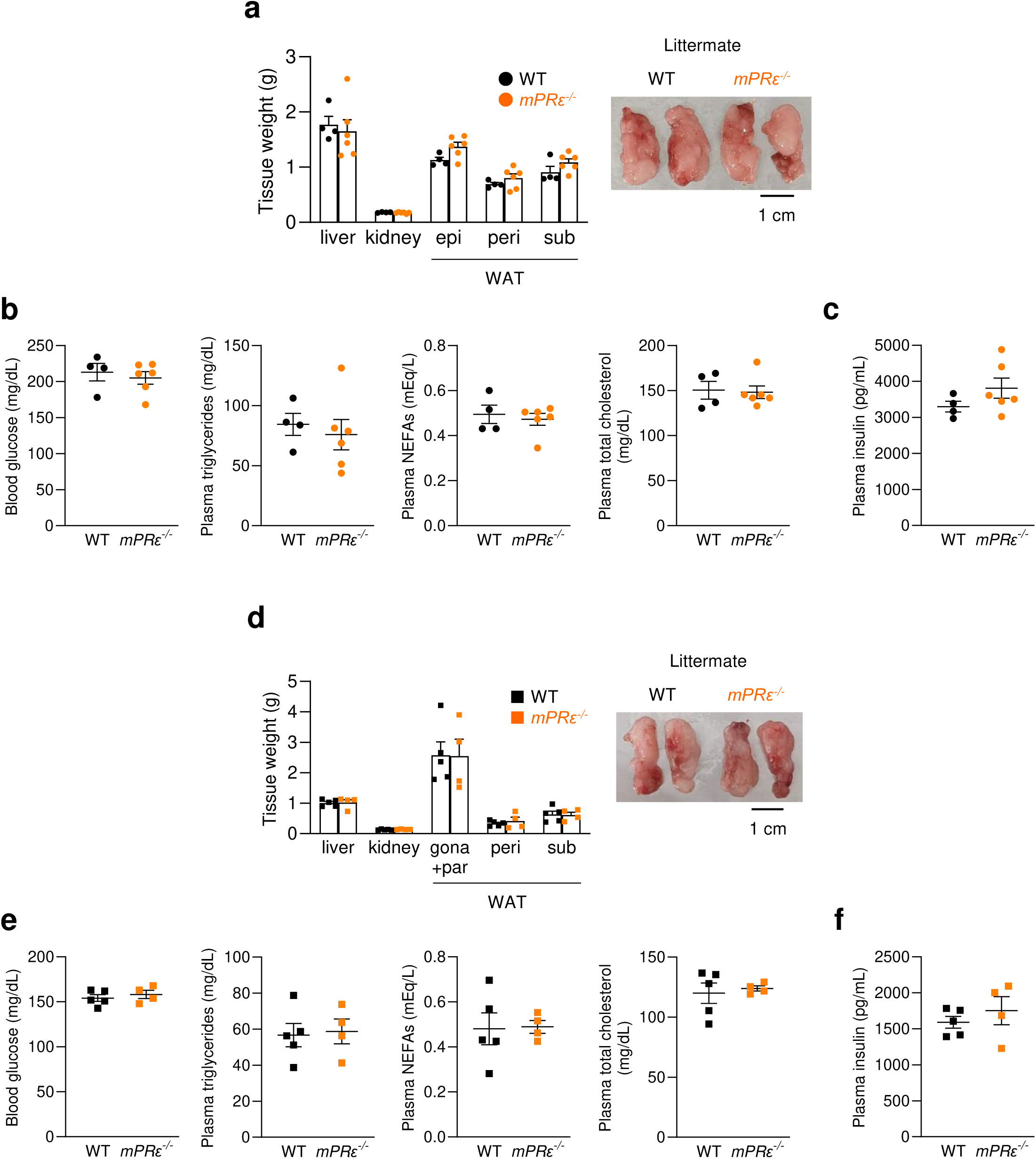
Metabolic parameters in *mPRε^-/-^* mice from *mPRε^+/-^* male and *mPRε^+/-^* female mice. (a) The tissue weight after 12 weeks in *mPRε^-/-^* male mice under high-fat diet (HFD) feeding (n = 4–6 from four litters; independent experiments). Epi, epididymal; peri, perirenal; sub, subcutaneous; BAT, brown adipose tissue; WATs, white adipose tissues. (b) Blood glucose, plasma triglyceride, non-esterified fatty acids (NEFAs), and total cholesterol levels in WT and *mPRε^-/-^* male mice after HFD intervention for 12 weeks (n = 4–6). (c) Plasma insulin levels in male mice (n = 4–6 from four litters; independent experiments). (d) The tissue weight after 12 weeks in *mPRε^-/-^* female mice under HFD feeding (n = 4–5 from four litters; independent experiments). Gona, gonadal; par, parametrial. (e) Blood glucose, plasma triglyceride, NEFAs, and total cholesterol levels in WT and *mPRε^-/-^* female mice after HFD intervention for 12 weeks (n = 4–5). (f) Plasma insulin levels in female mice (n = 4–5 from four litters; independent experiments). **P < 0.01, *P < 0.05 (Mann–Whitney U test: A; two-way ANOVA with the Bonferroni: B; Student’s t test: c–f). All data are presented as the mean ± SEM.

**Extended Data Fig. 3.**
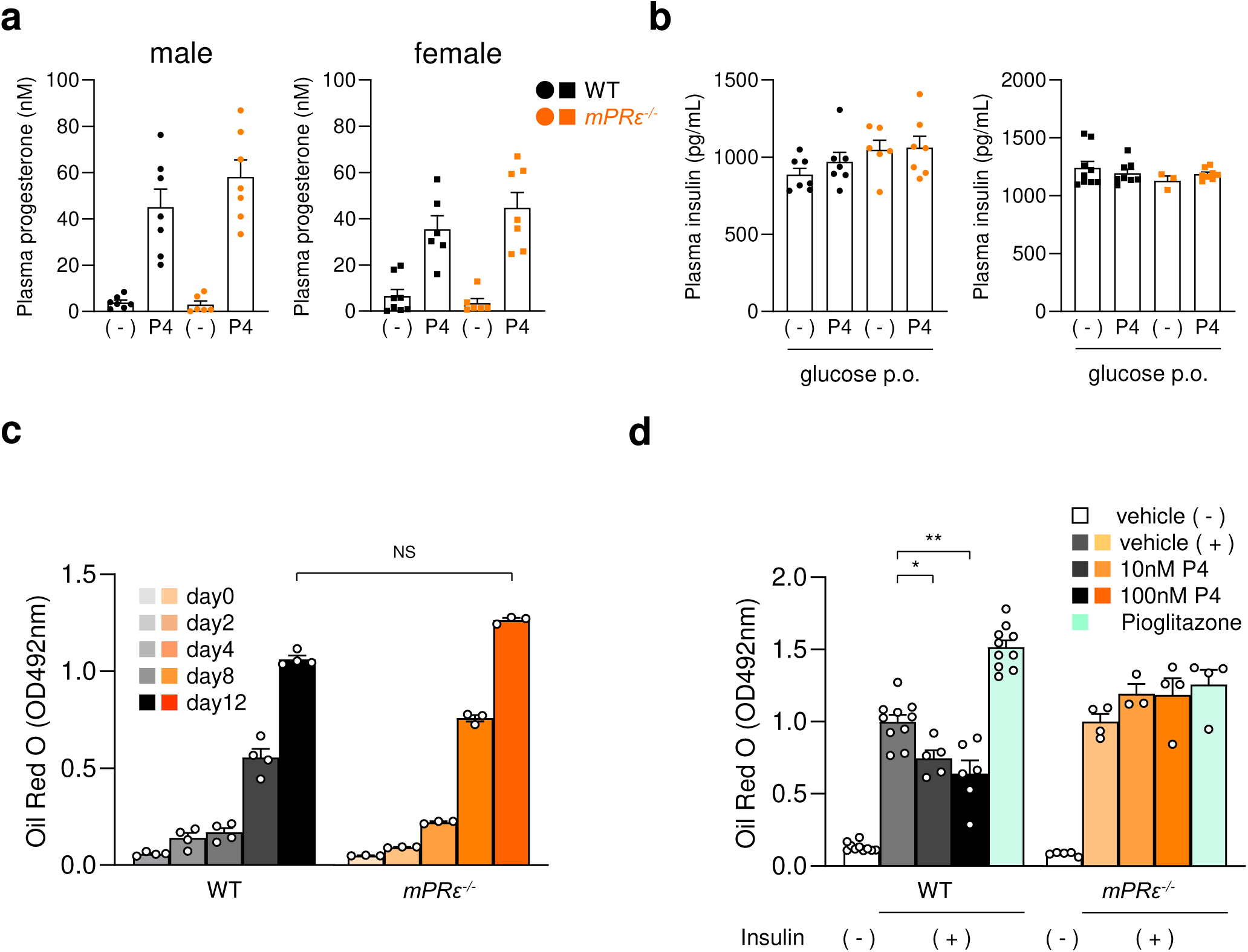
Functional analysis of adipose mPRε on glucose homeostasis. (a) Plasma progesterone levels 30 min after progesterone administration (s.c.) in male (left) and female (right) mice (n = 6–8). (b) Plasma insulin levels 30 min after administration of progesterone (s.c.) in male (left) and female (right) mice (n = 6–9). (c) Oil Red O staining of differentiated adipocytes derived from *mPRε^-/-^* mouse embryonic fibroblasts (MEFs) treated with 10 μM pioglitazone and inducer during adipogenesis (n = 3–4). (d) Oil Red O staining of MEF-derived adipocytes of *mPRε^-/-^* mice with progesterone addition (n = 3–10). **P < 0.01, *P < 0.05 (Dunn’s test). Results are presented as the mean ± standard error of the mean (SE). N.S.: not significant.

**Extended Data Fig. 4.**
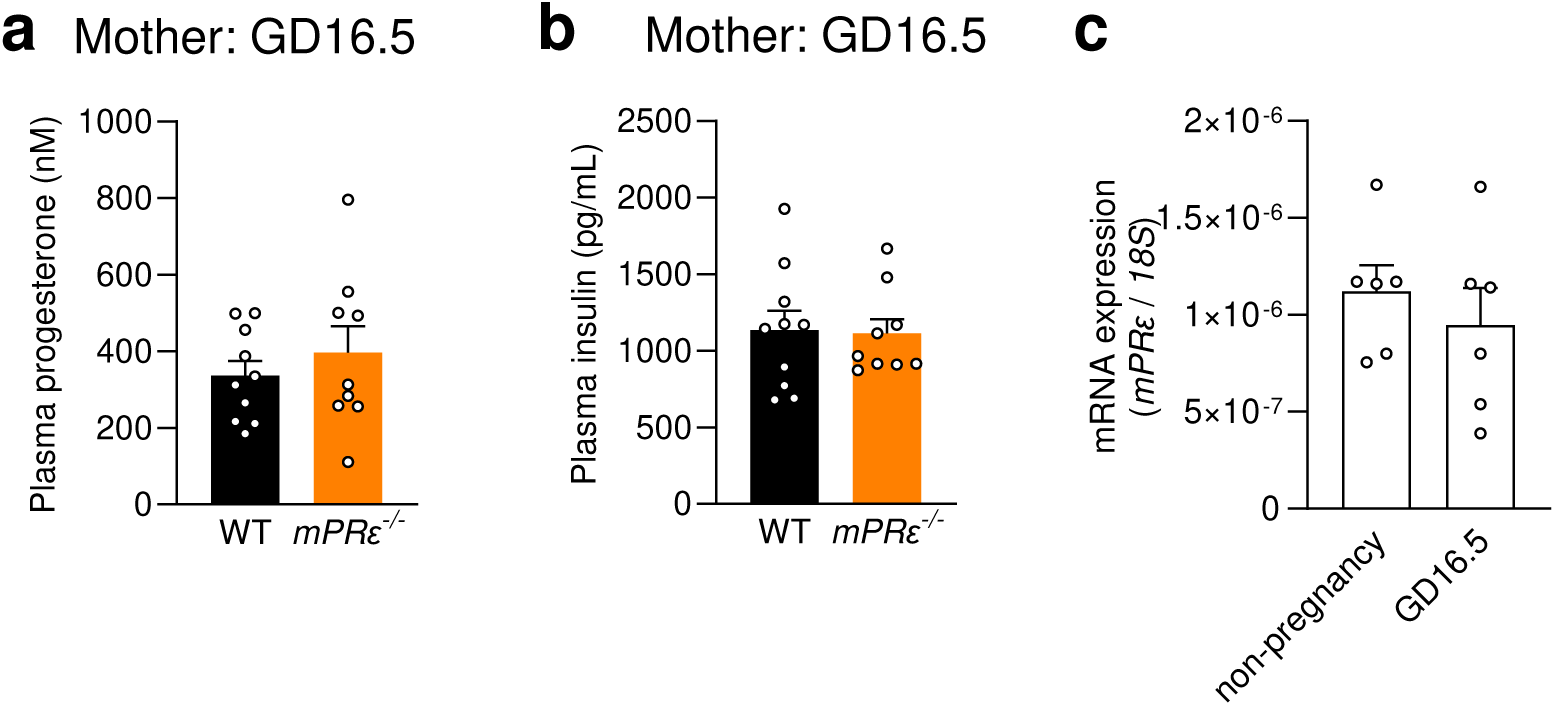
Functional analysis of maternal adipose mPRε in pregnancy. (a) Plasma progesterone levels in GD16.5 mothers (n = 9–11). (b) Plasma insulin levels in GD16.5 mothers (n = 9–11). (c) Comparison of adipose *mPRε* expression between non-pregnant mice and GD16.5 mothers (n = 6). Results are presented as the mean ± standard error of the mean (SE).

**Extended Data Fig. 5.**
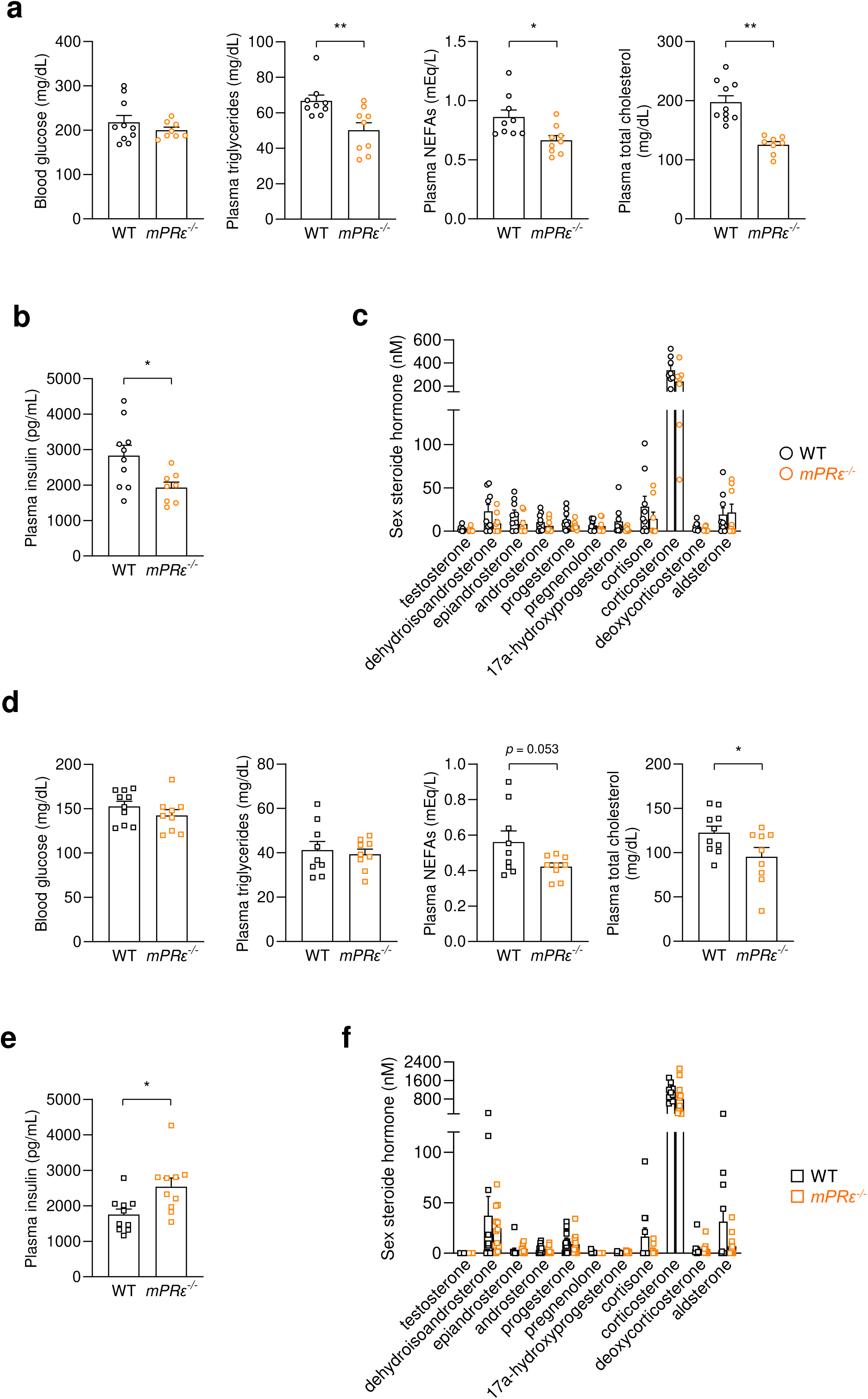
Metabolic parameters in *mPRε^-/-^* mice from *mPRε^-/-^* male and *mPRε^-/-^* female mice. (a) Blood glucose, plasma triglyceride, non-esterified fatty acids (NEFAs), and total cholesterols levels in WT and *mPRε^-/-^*male mice after HFD intervention for 12 weeks (n = 9–10). (b) Plasma insulin levels in male mice (n = 8–10 from three litters; independent experiments). (c) Plasma steroid levels at 16 weeks of age in male mice (n = 8–9). (d) Blood glucose, plasma triglyceride, NEFAs, and total cholesterol levels in WT and *mPRε^-/-^* female mice after HFD intervention for 12 weeks (n = 9–10). (e) Plasma insulin levels in female mice (n = 10 from three litters; independent experiments). (e) Plasma steroid levels at 16 weeks of age in female mice (n = 11–12). **P < 0.01, *P < 0.05 (Mann–Whitney U test: A; two-way ANOVA with the Bonferroni: B; Student’s t test: c–f). All data are presented as the mean ± SEM.

**Extended Data Fig. 6.**
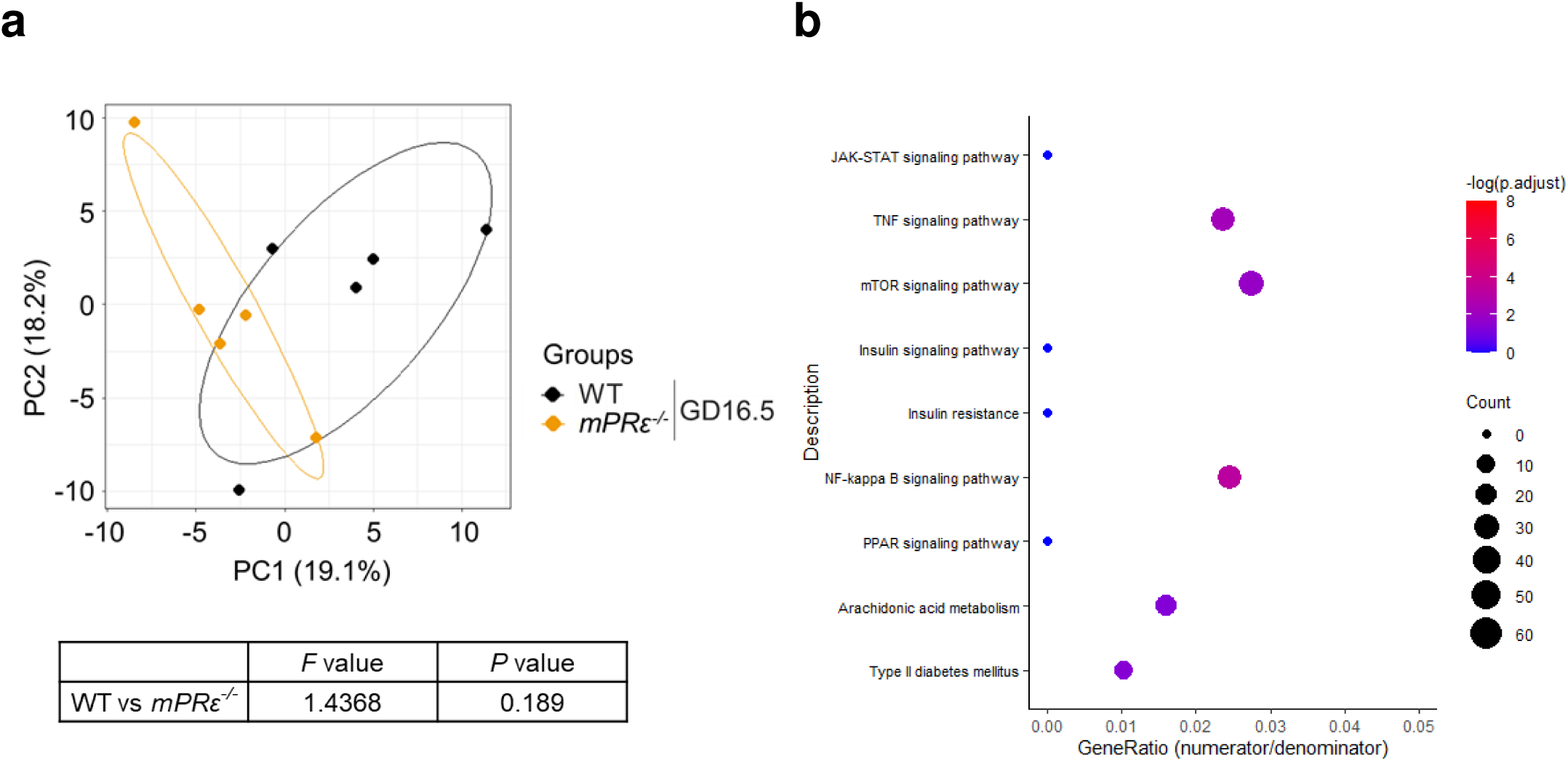
RNA sequencing in the liver of pregnant *mPRε^-/-^*mice. (a) The beta diversity as shown via principal component analysis (PCA) based on genes from KEGG (ID mmu00590; mmu04064; mmu04910) in the liver of WT and *mPRε^-/-^* mice with GD16.5 (n = 5). (b) KEGG enrichment analysis related to molecular function in the liver of GD16.5 *mPRε^-/-^* mother. P-values were adjusted based on the false discovery rate (FDR).

**Extended Data Fig. 7.**
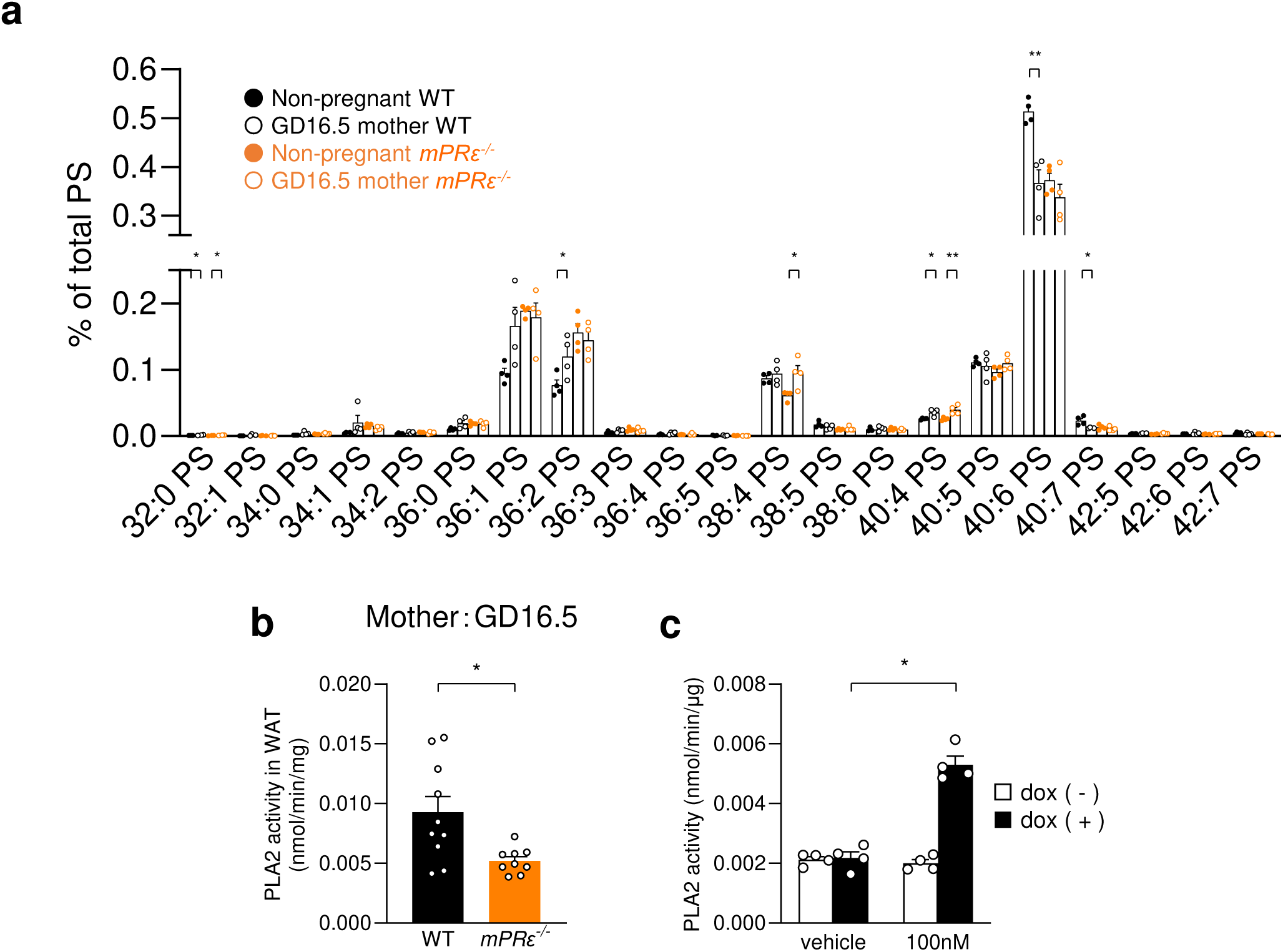
Progesterone-stimulated mPRε activation promotes PLA_2_ activity. (a) Membrane phospholipids of phosphatidylserine were quantified in the WAT of non-pregnant mice or GD16.5 mothers (n = 4 per group). (b) PLA_2_ activity in WAT of *mPRε^-/-^* GD16.5 mothers (n = 9–10). (c) PLA_2_ activity in response to progesterone (100 nM) in Flp-In mPRε T-REx HEK293 cells treated with or without doxycycline (n = 4). Dox; doxycycline. **P < 0.01, *P < 0.05 (Mann–Whitney U test). The results are presented as the mean ± standard error of the mean (SE).

**Extended Data Fig. 8.**
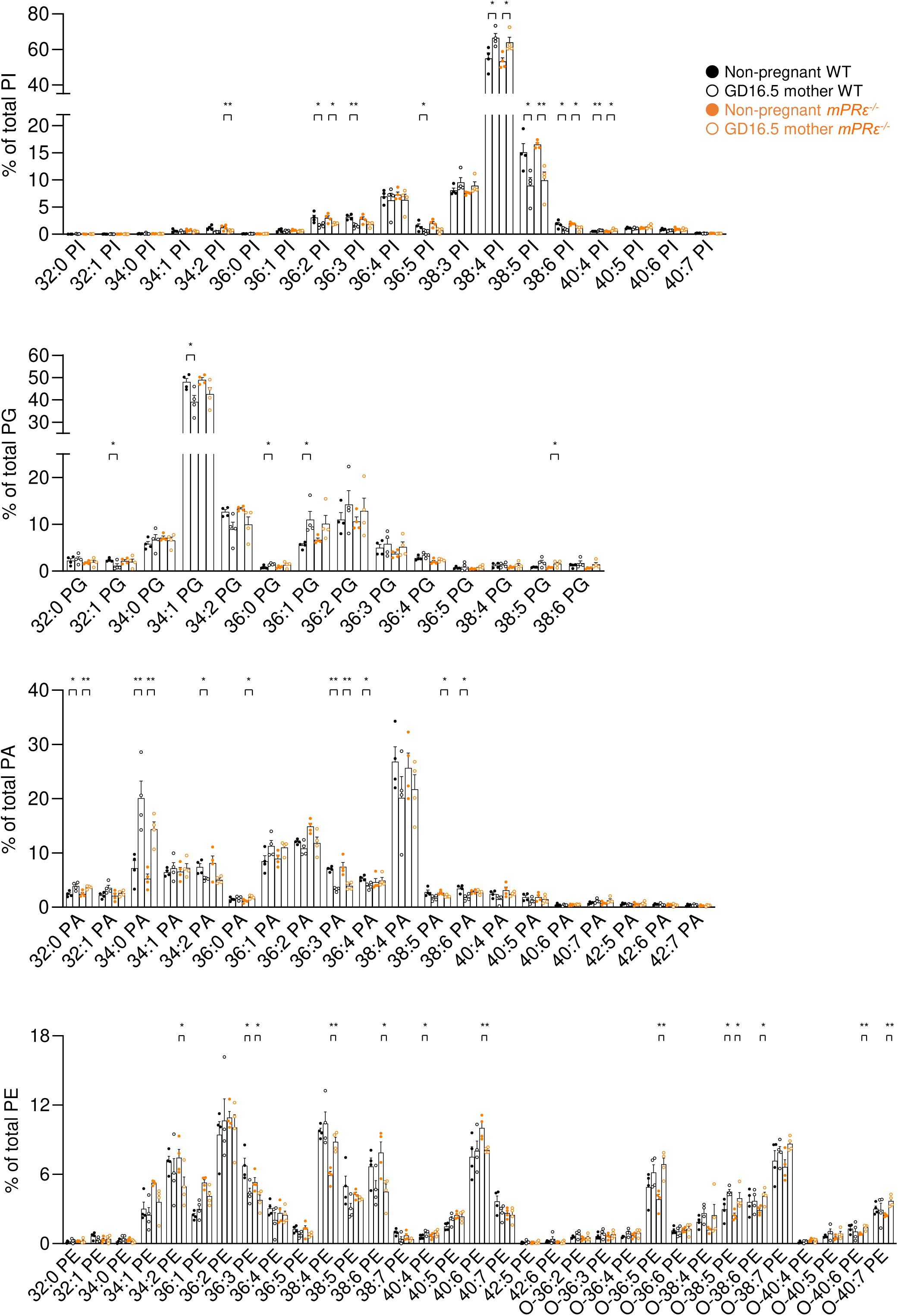
Phospholipids analysis in WAT of pregnant *mPRε^-/-^* mice. Quantitation of membrane phospholipids in WAT of GD16.5 *mPRε^-/-^* mother (n = 4). All data are presented as the mean ± SEM.

**Extended Data Fig. 9.**
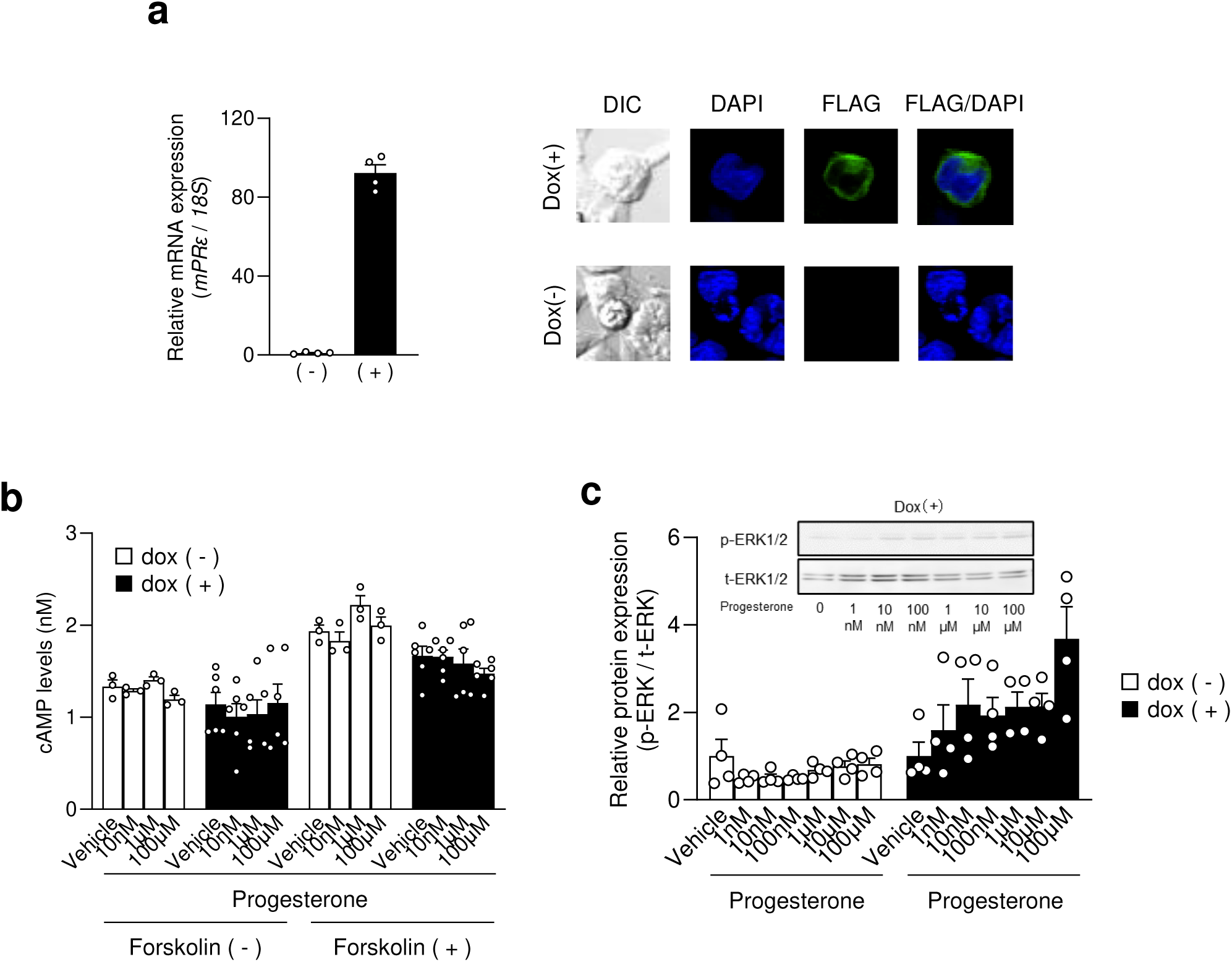
mPRε overexpressing HEK293 cells. (a) The mRNA expression of mPRε gene in Flp-In mPRε T-REx HEK293 cells treated with or without doxycycline (n = 4). (b) cAMP levels (n = 3–6) and (c) ERK1/2 phosphorylation (n = 4) in response to progesterone in a dose-dependent manner in Flp-In mPRε T-REx HEK293 cells treated with or without doxycycline. Intracellular cAMP levels were determined using a cAMP assay kit, and each data point is presented relative to the forskolin-induced cAMP levels. Dox; doxycycline. All data are presented as means ± SEM.

